# Among family variation in survival and gene expression uncovers adaptive genetic variation in a threatened fish

**DOI:** 10.1101/731497

**Authors:** Avril M. Harder, Janna R. Willoughby, William R. Ardren, Mark R. Christie

## Abstract

Variation in among-family transcriptional responses to different environmental conditions can help to identify adaptive genetic variation, even prior to a selective event. Coupling differential gene expression with formal survival analyses allows for the disentanglement of treatment effects, required for understanding how individuals plastically respond to environmental stressors, from the adaptive genetic variation responsible for among-family variation in survival and gene expression. We applied this experimental design to investigate responses to an emerging conservation issue, thiamine (vitamin B_1_) deficiency, in a threatened population of Atlantic salmon (*Salmo salar*). Thiamine is an essential vitamin that is increasingly limited in many ecosystems. In Lake Champlain, Atlantic salmon cannot acquire thiamine in sufficient quantities to support natural reproduction; fertilized eggs must be reared in hatcheries and treated with supplemental thiamine. We evaluated transcriptional responses (RNA-seq) to thiamine treatment across families and found 3,616 genes differentially expressed between control (no supplemental thiamine) and treatment individuals. Fewer genes changed expression additively (i.e., equally among families) than non-additively (i.e., family-by-treatment effects) in response to thiamine. Differentially expressed genes were related to known physiological effects of thiamine deficiency, including oxidative stress, cardiovascular irregularities, and neurological abnormalities. We also identified 1,446 putatively adaptive genes that were strongly associated with among-family survival in the absence of thiamine treatment, many of which related to neurogenesis and visual perception. Our results highlight the utility of coupling RNA-seq with formal survival analyses to identify candidate genes that underlie the among-family variation in survival required for an adaptive response to natural selection.

## Introduction

Understanding if and how species can adapt to rapidly changing environmental conditions is a primary goal of modern conservation biology (Bernatchez, 2016; Stockwell, Hendry, & Kinnison, 2003). One of the key challenges in meeting this goal is uncovering the adaptive genetic variation required for a response to selection and deciphering whether this adaptive genetic variation will be sufficient to respond to anthropogenically-induced agents of selection. Contemporary genomic approaches have revolutionized our ability to identify regions of the genome responding to selection, even over relatively short time periods (Franks, Kane, O’Hara, Tittes, & Rest, 2016; van’t Hof et al., 2016; Willoughby, Harder, Tennessen, Scribner, & Christie, 2018). However, such methods often lack sufficient power to detect rapid responses to selection, especially when examining polygenic traits shaped by large numbers of loci of small effect (Pritchard, Pickrell, & Coop, 2010; Wellenreuther & Hansson, 2016). Furthermore, genomic approaches can only provide insights after selection has already occurred; thus their utility for predicting responses to selection requires appropriate study systems or long-term experimental breeding designs. One alternative to these approaches is experimental transcriptomics. By carefully designing treatments, rearing F1 offspring in a common environment, and deeply sequencing mRNA, it is possible to uncover an adaptive, genetic response to selection (Christie, Marine, Fox, French, & Blouin, 2016; Passow et al., 2017; Uusi-Heikkilä, Sävilammi, Leder, Arlinghaus, & Primmer, 2017). Coupling family-level replication and formal survival analyses allows for the disentanglement of treatment effects, required for understanding how individuals plastically respond to environmental stressors, and among-family variation in survival and gene expression. Here, we apply these techniques to a threatened fish population, whose successful reintroduction will require an adaptive response to an emerging conservation issue, thiamine deficiency.

Evidence is mounting that populations of diverse taxa are becoming increasingly deficient in thiamine (vitamin B_1_) (Balk et al., 2009, 2016). For example, high rates of mortality or reduced reproductive success associated with thiamine deficiency have been observed in invertebrates (Balk et al., 2016), fishes (Futia et al., 2017), reptiles (Honeyfield et al., 2008; Ross et al., 2009), and birds (Balk et al., 2009). Furthermore, many cases of thiamine deficiency remain undetected. From a conservation standpoint, it is particularly concerning that thiamine deficiency remains largely undetected despite potentially being a large driver of population declines.

Thiamine is an essential vitamin that is synthesized by prokaryotes, plants, and fungi; animals are incapable of producing thiamine and primarily acquire the vitamin through their diets (Bettendorff, 2013). The physiological manifestations of thiamine deficiency are directly related to thiamine’s roles in bioenergetic, neurological, and cardiovascular pathways. Thiamine serves as a cofactor for enzymes in metabolism and energy production pathways (*i.e*., pentose phosphate pathway and tricarboxylic acid cycle) and thiamine deficiency leads to extreme lethargy (Brown et al., 2005; Fitzsimons, Brown, Honeyfield, & Hnath, 1999). Thiamine is also required for production of neurotransmitters, antioxidants, and myelin (Bettendorff, 2013), consistent with the neurological and behavioral signs of thiamine deficiency, including brain lesions (Butterworth, 2009; Honeyfield et al., 2008; Lee, Jaroszewska, Dabrowski, Czesny, & Rinchard, 2009) and uncoordinated movements (Brown et al., 2005; Fisher, Spitzbergen, et al., 1995; Fitzsimons et al., 2005; Sechi & Serra, 2007). Thiamine deficiency can also impair cardiovascular function, leading to low blood pressure, irregular heart rate, pulmonary edema, and circulatory collapse (Essa et al., 2011; Sechi & Serra, 2007). Because thiamine plays a central role in growth, development, and proper neurological function (Bettendorff, 2013), thiamine deficiency can impair an individual’s capacity to forage, avoid predation, and reproduce (Carvalho et al., 2009; Fisher, Spitzbergen, et al., 1995; Fitzsimons et al., 2009), all of which can contribute to large reductions in population size (Ketola, Bowser, Wooster, Wedge, & Hurst, 2000; Mörner et al., 2017).

The underlying causes of thiamine deficiency vary among taxa and environments. In fishes, the emergence of thiamine deficiency is largely attributed to diet. For example, thiamine deficiency has often been observed in salmonids with diets containing alewife (*Alosa pseudoharengus*) and rainbow smelt (*Osmerus mordax*), both of which contain high levels of thiaminase, a thiamine-degrading enzyme (reviewed in Harder et al., 2018). In the Baltic Sea, the occurrence of thiamine deficiency in Atlantic salmon (*Salmo salar*) also coincides with the consumption of fishes with low thiamine:fat content ratios, including Atlantic herring (*Clupea harengus*) and sprat (*Sprattus sprattus*) (Hansson et al., 2001; Keinänen et al., 2012). However, these fishes also contain thiaminase, making it difficult to establish low thiamine:fat content ratios as direct, causative agents of thiamine deficiency. For adult salmon returning to spawn, the most obvious signs of thiamine deficiency are uncoordinated, “wiggling” swimming patterns and an inability to remain upright in the water column (Fisher, Spitsbergen, Iamonte, Little, & Delonay, 1995; Fitzsimons et al., 2005). If thiamine deficient individuals are able to spawn, these behaviors are inevitably mirrored in their offspring. Individuals hatching from thiamine deficient eggs do not survive for more than a few weeks and exhibit physical signs of deficiency, such as hemorrhaging and large yolk sacs with opacities and edema, prior to death (Fig. 1A) (Fisher, Spitsbergen, et al., 1995). The inability of thiamine deficient salmon to successfully reproduce is an emerging conservation and management issue (reviewed in Harder et al., 2018), and impedes reintroduction efforts throughout their native range.

**Figure 1.**
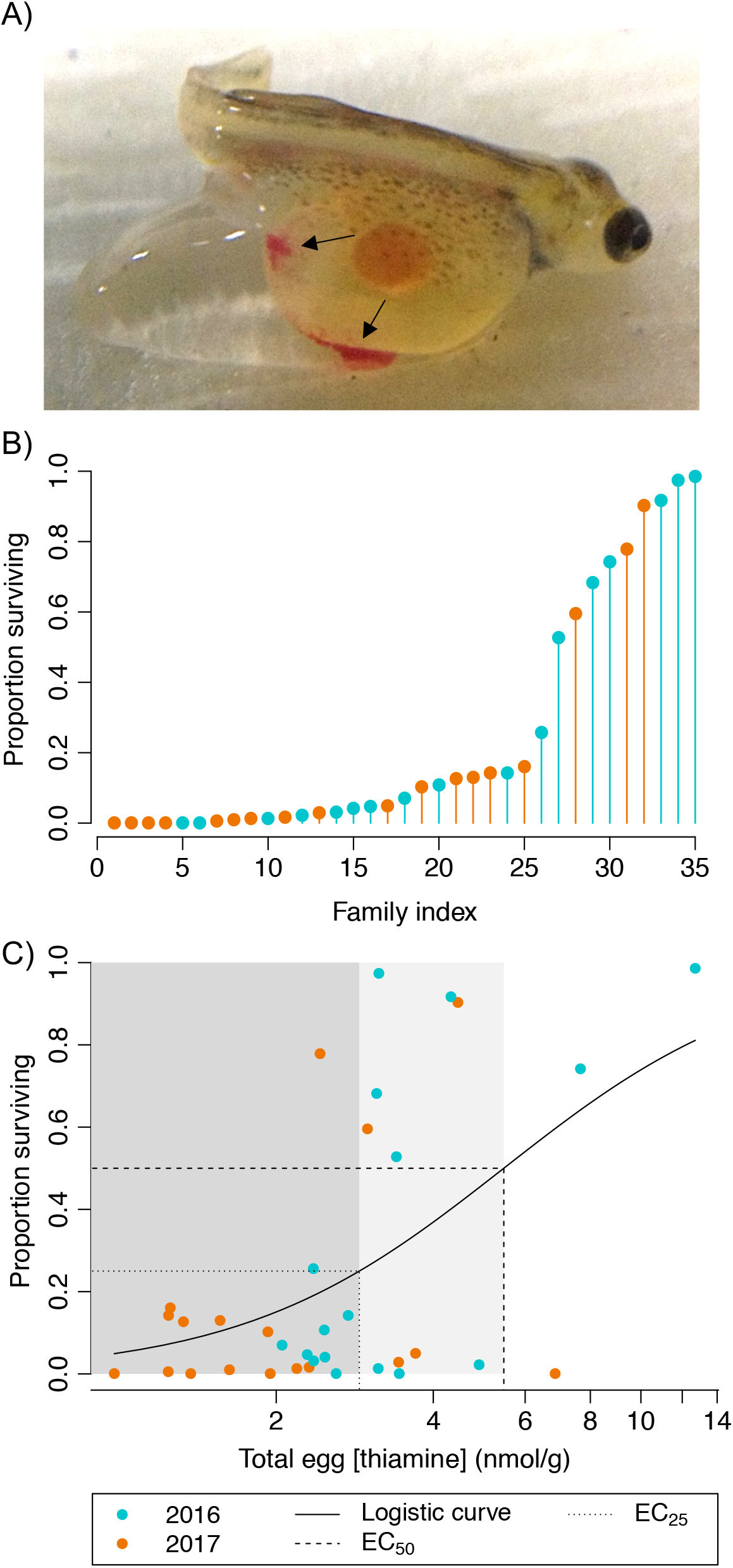
A) Atlantic salmon fry exhibiting characteristic signs of thiamine deficiency, including hemorrhaging (indicated by arrows) and edema in the posterior portion of the yolk sac. B) Proportion of untreated fry surviving to the onset of exogenous feeding for 35 families spawned in 2016 and 2017. C) Dose-response curve illustrating the relationship between total egg thiamine concentration (nmol/g) and proportion of untreated fry surviving to yolk sac absorption. Shaded grey areas highlight families with egg [thiamine] < EC_25_ (dark grey) and with egg [thiamine] >EC_25_ but <EC_50_ (light grey), where EC_25_ and EC_50_ equal the effective concentrations required for 25% and 50% survival, respectively.

One such reintroduction effort occurs in Lake Champlain (Canada and USA), where Atlantic salmon were extirpated from the lake in the early 1800s (Marsden & Langdon, 2012; SI Introduction). Diversifying the forage base or controlling the alewife population in Lake Champlain could alleviate thiamine deficiency in Atlantic salmon, but efforts to eradicate invasive species after population establishment are often prohibitively expensive and the possibility of reinvasion cannot be eliminated (Myers, Simberloff, Kuris, & Carey, 2000). Alternatively, recent research suggests that Atlantic salmon populations with diets high in thiaminase may have genetically adapted to low thiamine availability (Houde, Saez, Wilson, Bureau, & Neff, 2015). This rapid genetic adaptation could be the result of selection on genes associated with thiamine-dependent pathways. For example, conformational changes in enzymes requiring thiamine as a cofactor could increase the binding affinity for thiamine or, alternatively, variation in regulatory sequences could modify the expression of genes involved in thiamine uptake and intracellular transport. However, the application of supplemental thiamine to all fertilized eggs reared in the Lake Champlain hatchery precludes selection related to thiamine deficiency at early life stages, and it is currently unknown whether genetic variation in this population could support a response to such selection. By coupling thiamine treatments, RNA-seq, and survival analyses on F1 offspring from 9 families raised in a common environment, we identified an among-family adaptive response in a thiamine-deficient population of Atlantic salmon and identified pathways and functions impacted by thiamine deficiency. Categorizing relationships between gene expression and survivorship patterns revealed two distinct groups of differentially expressed genes that (1) underlie putatively adaptive responses to thiamine deficiency among families and (2) reflect the treatment effect of thiamine use regardless of genetic differences. Our results are consistent with a heritable, among-family basis for tolerance to low thiamine availability.

## Methods

### Study system and experimental crosses

We collected gametes from 35 pairs of adult male and female Atlantic salmon returning to the Ed Weed Fish Culture Station (Grand Isle, Vermont, USA) across two spawning seasons: 17 pairs in November 2016 and 18 pairs in November 2017. We immediately froze approximately 50 eggs from each female on dry ice for total thiamine concentration analysis whereby two, 1-g biological replicates of unfertilized egg tissue were analyzed via high performance liquid chromatography (*sensu stricto* Futia et al. 2017). We also performed total thiamine concentration analyses on 2-g samples of maternal muscle tissue sampled from each female (two samples per female) during U.S. Fish and Wildlife Service disease testing procedures. We transported gametes at 4 °C to the White River National Fish Hatchery (Bethel, Vermont, USA), where we systematically combined milt and eggs to generate 35 families (see SI Methods for crossing details). We divided fertilized eggs from each family into two groups, placing one group into a 1% thiamine mononitrate solution (hereafter, “treated”) and the other into a control water bath (hereafter, “untreated”). After 30 minutes, we rinsed all eggs with fresh water and transferred them to heath trays with one tray per family and treatment combination. We left the eggs undisturbed until reaching the eyed stage (when individuals exhibit retinal pigmentation, approximately 50 days post fertilization), at which point we counted and removed inviable eggs. After the eyed stage was reached, we recorded mortality and removed inviable eggs from all trays each week. Hatching occurred approximately 75 days post fertilization and we continued to monitor and remove dead individuals from all trays each week for the remainder of the experiments. We concluded the experiments after surviving fry had absorbed their yolk sacs (~130 days post fertilization) and just prior to initiation of exogenous feeding.

### Sampling for RNA-seq

At 95 days post fertilization, we sampled a total of thirty-six individuals for gene expression analyses from 9 of the 18 families spawned in 2017. Due to hatchery broodstock quotas for treated individuals, we were limited to sampling these 9 families. To control for variation in development, we only sampled from families that were spawned on the same day. We froze individuals from each family and treatment group in dry cryogenic shipping dewars charged with liquid nitrogen and shipped them to Purdue University for storage at −80 °C. We subsequently placed two frozen individuals from each family and treatment group (n = 9 families * 2 treatments (+/− supplemental thiamine) * 2 individuals = 36) into 10 volumes of RNAlater-ICE (Invitrogen) pre-chilled to −80 °C and allowed the samples to reach −20 °C overnight. We then homogenized samples using a TissueRuptor II (Qiagen) and extracted total RNA from each homogenate using an RNeasy kit (Qiagen).

### Survival analyses

We generated a dose-response curve for egg thiamine concentration and proportion of untreated individuals in each family surviving at the end of the experiments with the *drc* package (Ritz, Baty, Streibig, & Gerhard, 2015) in R version 3.5.3 (R Core Team, 2019). We selected the appropriate model by using the mselect function to calculate AIC values, with the two-parameter log-logistic function having the lowest AIC value. We next calculated effective concentrations of egg thiamine required for 25% and 50% survival (EC25 and EC50, respectively) from the resulting logistic curve.

For the 9 families used in RNA-seq, we conducted survival analyses to determine whether treatment affected survival within a family over time and to determine the relative risks of death associated with belonging to each family according to survivorship of untreated individuals. We constructed Kaplan-Meier survival distributions for each family and treatment combination and used a log-rank test to determine whether treatment status significantly affected survival within each family (Kleinbaum & Klein, 2012). We then compared survival distributions for untreated individuals from each family against the survival distribution for untreated individuals from family A (the family with the highest survival rate of all families). We used Cox proportional hazards regressions to calculate hazard ratio values for all families (Cox, 1972) using the *survival* package (Therneau, 2015) in R. We censored individuals removed for RNA-seq in the analysis. For each family, the calculated hazard ratio represents the probability of mortality associated with belonging to that family, compared to family A. We also conducted a linear regression to test for a relationship between hazard ratio value and egg thiamine concentration. To meet assumptions of normality, we log-transformed hazard ratio values prior to all regression analyses.

When spawning families for this study, reciprocal crosses were not feasible due to limited egg and milt availability, therefore, we could not formally test for maternal effects (*sensu* Christie et al., 2016). However, female size is often correlated with offspring size, and larger offspring frequently exhibit higher fitness than smaller offspring in a common environment (reviewed in Marshall, Heppell, Munch, & Warner, 2010). We therefore performed linear regressions to test for relationships between maternal physical characteristics (*i.e*., standard length and weight) and proportion of untreated offspring surviving at the end of the experiment. We also plotted maternal muscle and egg thiamine concentration against proportion of untreated offspring surviving for the 9 families sampled for RNA-seq. A strong association between maternal characteristics and untreated offspring survival might indicate that maternal condition plays a larger role in determining thiamine deficiency outcomes than among-family genetic variation.

### RNA-seq and sequence processing

We assessed total RNA concentration and quality on an Agilent BioAnalyzer at the Purdue Genomics Core Facility, with sample RIN scores ranging from 9.3-10.0. One library was prepared for each individual using the TruSeq Stranded mRNA protocol (Illumina) and cDNA was sequenced on an Illumina NovaSeq 6000 to generate an average of 87 million 150 bp paired-end reads per library (Table S1). We removed adapter sequences and clipped poor quality bases (quality score < 20) from both ends of reads using Trimmomatic (Bolger, Lohse, & Usadel, 2014) and aligned reads to the annotated Atlantic salmon reference genome (*S. salar* ICSASG_v2 assembly, NCBI accession GCA_000233375.4; Lien et al., 2016) using HISAT2 (Kim, Langmead, & Salzberg, 2015) with the --*downstream-transcriptome-assembly* option and reporting primary alignments. We next assembled transcripts for each sample using StringTie (Pertea et al., 2015) default parameters and the Atlantic salmon reference annotation file (ICSASG_v2) to guide assembly, and merged sample transcripts using StringTie. A transcript count matrix was next created with featureCounts (Liao, Smyth, & Shi, 2014), excluding chimeric fragments (-*C* option) and requiring that both reads in a pair be successfully mapped (-*B* option). By default, featureCounts does not count reads with multiple alignments (*i.e*., a single read aligned to multiple locations in the reference) or read pairs that overlap multiple features, and we retained these settings in our analyses.

### Differential expression analyses: treatment effects

We first made comparisons between treated and untreated individuals using both an *a priori* list of reference genes and a standard discovery-based gene identification pipeline. We generated a list of *a priori* genes predicted to be differentially expressed between treated and untreated samples using 4 criteria: (1) genes associated with thiamine-related biological process gene ontology (GO) terms (any line containing “thiamine” in Ssal_ICSASG_v2_GOAccession.txt downloaded from SalmoBase (Samy et al., 2017) on June 28, 2018), (2) genes encoding thiamine diphosphate (TDP) dependent enzymes, (3) genes encoding enzymes that contain a TDP binding site (NCBI conserved protein domain family “TPP_enzymes”), and (4) genes included in the *S. salar* thiamine metabolism pathway in the NCBI BioSystems Database (BSID: 1429556).

We conducted differential gene expression analyses separately in DESeq2 (Love, Huber, & Anders, 2014) for: 1. the *a priori* list of predicted differentially expressed genes (DEGs) and 2. the list of all assembled transcripts. We identified DEGs associated with thiamine treatment status while controlling for the effects of family, and considered genes with an FDR-adjusted *p*-value (*p*_adj_) < 0.05 to be differentially expressed. We used the *prcomp* command in R to conduct a principal component analysis for DEGs identified from the list of all assembled transcripts.

Using the count matrix for all samples, we identified modules of co-expressed genes by calculating pair-wise Pearson correlations between each pair of genes using the WGCNA package (Langfelder & Horvath, 2008). We set the minimum modules size to 30 genes and merged correlated modules (*r^2^* > 0.9). Each module comprised genes that showed similar expression patterns across samples within a treatment. Following the approach outlined in Langfelder and Horvath (2008) we performed the following steps. First, we summarized module expression using a principal components analysis (PCA) and calculated eigengenes as the first principal component (PC1) for each module. Second, we used the Pearson correlation to search for associations between module eigengenes and treatment status, and calculated *p*-values for correlations using a Student’s asymptotic test. Finally, we applied a Bonferroni correction to account for multiple testing.

For each module significantly associated with treatment status, we performed a gene ontology (GO) enrichment analysis to identify which Biological Process GO terms associated with the DEGs were overrepresented compared to the genome-wide complement of *S. salar* GO terms (*p* < 0.001). We used the TopGo package in R (Alexa & Rahnenfuhrer, 2016), which is less biased towards the most general GO terms because it employs a hierarchical methodology, and chose the ‘weight01’ algorithm because this method efficiently identifies enriched terms at all levels of the GO hierarchy while limiting the proportion of false positives (Alexa, Rahnenfuhrer, & Lengauer, 2006). After identifying overrepresented GO terms in each module, we created a list of terms unique to each module (all overrepresented terms shared among all 3 modules are provided in Table S2). For each module, we created a list of the top 20 genes ranked by gene significance (a value calculated in WGCNA that indicates the biological significance of a module gene with respect to the explanatory variable of interest). We used unique GO terms associated with the top 20 genes to construct a network of GO terms for each module, and the *metacoder* package (Foster, Sharpton, & Grünwald, 2017) to visualize networks in R. We pruned internal nodes from each network for ease of visualization.

### Identifying putatively adaptive genes

To identify putatively adaptive genes that could respond to selection imposed by thiamine deficiency, we generated a transcript count matrix for untreated individuals only. We conducted differential gene expression analysis on this group in DESeq2 in R with family hazard ratio value as the explanatory variable. We considered genes with *p*_adj_ < 0.05 and with a fold-change > 1 (log_2_ fold-change > 0.5 between the families with the lowest and highest hazard ratio values) to be putatively adaptive.

We further categorized the adaptively expressed genes by whether increasing hazard ratio (*i.e*., increasing probability of mortality) was associated with either an increase or a decrease in gene expression, when analyzed across families. We further filtered genes belonging to each category by applying a linear regression approach to each gene, with log(hazard ratio) as the explanatory variable and overall gene expression (fragments per million mapped fragments, FPM) as the response variable. To account for the fact that we sequenced two siblings from each family, we conducted each regression using 1 randomly selected individual from each family, and repeated this process 1,000 times per gene. We calculated coefficient means for each gene and variances in the means as 95% confidence intervals. We discarded genes from further analyses if the slope of the regression did not differ from 0 or if the adjusted r^2^ of the regression was < 0.3. For each group of putatively adaptive genes, we performed a gene ontology (GO) enrichment analysis using the same approach described above. We ranked GO terms by *p*-value for each category and retained the top 50 terms from each group (*p* < 0.001 for all retained terms).

### Categorizing treatment effects: additive vs. family x treatment interactions

To categorize treatment effects, we first limited our analyses to genes previously identified as differentially expressed with respect to thiamine treatment (see “Differential expression analyses: treatment effects” section above). Additive effects occur when the response to the thiamine treatment was equal across families. When the slopes of both treatment and controls do not differ from zero across families, this pattern represents a purely environmental response to thiamine treatment. By contrast, one family may respond to thiamine treatment differently than another, and this pattern can result in a family x treatment interaction. Using this approach, we can disentangle among-family (*i.e*., putatively adaptive) variation in gene expression from both an additive (purely environmental) response to treatment and a family x treatment interaction. We calculated regressions separately for each treatment group with log(hazard ratio) as the explanatory variable and fragments per million mapped fragments (FPM) as the response variable. We again conducted each regression using 1 randomly selected individual from each family and treatment combination and repeated this process 1,000 times per gene. We identified significant differences between the treatment and control groups by comparing the bootstrapped coefficient estimates for slope and intercept. To approximate a significance cut off of α = 0.05, we identified genes where the mean coefficient estimate +/− 1 standard error (approximated by 83% quantiles; Payton, Greenstone, & Schenker, 2003) between the treatment and control groups did not overlap. In addition, slopes were considered to not differ from zero if their 95% confidence intervals included zero. We also categorized genes according to whether or not the slopes or intercepts of the treated and untreated regression lines differed from one another.

## Results

### Thiamine concentration and survival analyses

In 2016 and 2017, the proportion of untreated individuals surviving within each family varied widely and ranged from 0 to 0.99 (mean = 0.25, SD = 0.34) (Fig. 1B). Total thiamine concentrations in unfertilized eggs were also variable and ranged from 0.98 to 12.71 nmol total thiamine/g unfertilized egg tissue (mean = 3.09 nmol/g, SD = 2.23 nmol/g). Fitting a dose) response curve to the relationship between egg thiamine concentration (nmol/g) and proportion of untreated individuals surviving at the end of each experiment resulted in an EC_25_ of 2.89 nmol/g and an EC50 of 5.46 nmol/g (*i.e*., 5.46 nmol/g of thiamine is required for 50% survival) (Fig. 1C).

Within each of the 9 families sampled for gene expression analyses, Kaplan-Meier survival distributions for treated and untreated individuals were significantly different (log-rank test; family A: *p* < 0.01, families B-I: *p* < 0.0001), indicating that thiamine treatment significantly and positively impacted survival over time for all families (Fig. S1). Hazard ratios ranged from 1 (for reference family A) to 80.12 (family I). Hazard ratio values > 1 indicate that a higher risk of death is associated with belonging to a particular family (*i.e*., the risk of death) associated with belonging to family I is 80.12 times greater than the risk of death associated with ‘ belonging to family A).

### Differential expression analyses: treatment effects

) Across all 36 individuals sequenced, the average rate of single concordant alignment for read pairs per sample was 80.9% and 62.2% of read pairs were successfully assigned to annotated features with featureCounts (Table S1). The final list of *a priori* genes included 106 unique genes. Of these genes, 17 were differentially expressed between treated and untreated individuals after controlling for false discovery (*p*_adj_ < 0.05; Table 1). Three of these genes—which encode adenylate kinase and reduced folate carrier—are involved in regulating intracellular concentrations of TDP (Fig. S2). Most of the remaining *a priori* DEGs comprise TDP-dependent enzymes or kinases that control TDP-dependent enzyme activity (Table 1). Differential expression analysis conducted using the full list of assembled transcripts resulted in the identification of 3,616 DEGs after controlling for false discovery (*p*_adj_ < 0.05; Table S3). A principal component analysis conducted with these DEGs showed treated samples clustering closely together, with PC1 differentiating treated and untreated individuals within each family and describing 59% of the variation (Fig. 2).

**Table 1.** Genes identified as differentially expressed that were hypothesized *a priori* to be implicated in thiamine deficiency. Gene symbols correspond to those used in *S. salar* NCBI assembly GCA_000233375.4 (Lien et al., 2016). Direction of log2 fold change values indicate direction of regulation in the treated group relative to the untreated group. Genes without chromosome arm information are located on unplaced scaffolds in the *S. salar* reference assembly.

| Gene symbol | Gene description | Chromosome arm | Log2 fold change | FDR corrected *p*-value |
| --- | --- | --- | --- | --- |
| bckdha | branched chain keto acid dehydrogenase E1, alpha polypeptide | 9qc | -0.166 | 1.17E-08 |
| LOC106600850 | pyruvate dehydrogenase (acetyl-transferring) kinase isozyme 2, mitochondrial-like | 3q | 0.476 | 6.09E-05 |
| kad | adenylate kinase 1-2 | 1qb | 0.185 | 6.44E-04 |
| kad | adenylate kinase | 11qb | 0.235 | 1.70E-03 |
| ilvbl | ilvB (bacterial acetolactate synthase)-like | 20qb | 0.165 | 1.70E-03 |
| LOC106581299 | alkaline phosphatase, tissue-nonspecific isozyme-like | 20qb | 0.638 | 1.70E-03 |
| LOC106583968 | DET1- and DDB1-associated protein 1-like | 23 | 0.260 | 1.70E-03 |
| LOC106569142 | 2-oxoglutarate dehydrogenase, mitochondrial-like | 1qb | -0.268 | 3.44E-03 |
| LOC106561658 | probable 2-oxoglutarate dehydrogenase E1 component DHKTD1, mitochondrial | 10qb | -0.288 | 1.63E-02 |
| LOC106587025 | 2-oxoglutarate dehydrogenase, mitochondrial | 26 | -0.226 | 1.63E-02 |
| hacl1 | 2-hydroxyacyl-CoA lyase 1 | -- | 0.241 | 2.20E-02 |
| LOC106579320 | pyruvate dehydrogenase (acetyl-transferring) kinase isozyme 2, mitochondrial-like | 19qb | -0.489 | 2.20E-02 |
| LOC106578452 | pyruvate dehydrogenase (acetyl-transferring) kinase isozyme 3, mitochondrial | 19qa | -0.214 | 2.28E-02 |
| LOC106580908 | thiamine transporter 2-like | 20qb | 0.969 | 2.28E-02 |
| LOC106560358 | thiamine transporter 2-like | 10qa | 1.862 | 2.85E-02 |
| LOC106563967 | dihydrolipoyllysine-residue acetyltransferase component of pyruvate dehydrogenase complex, mitochondrial-like | 11qb | -0.182 | 3.93E-02 |
| LOC106575607 | folate transporter 1-like | 17qa | 0.253 | 4.36E-02 |

**Figure 2.**
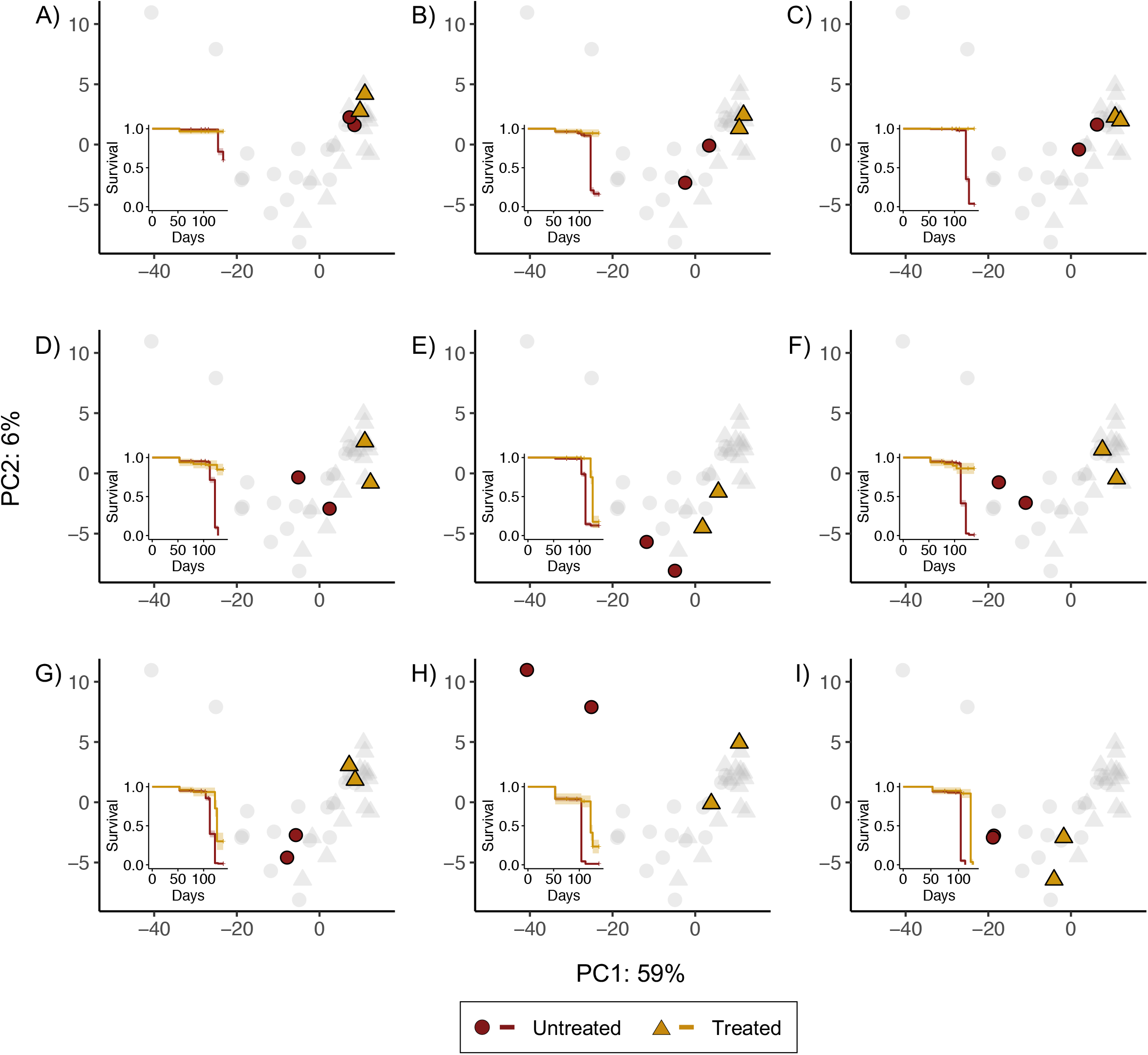
Principal component analysis performed using differentially expressed genes (n = 3,616). Results are presented by family (A-I), such that the four colored points plotted in each panel are full siblings. PC1 explains 59% of the variation and distinguishes among treated and untreated samples. Triangles represent treated individuals and circles represented untreated individuals. Insets are Kaplan-Meier survival distributions for treated (yellow) and untreated (red) individuals from each family. Inset *x*-axes represent time in days post fertilization, whereas the *y*-axes represent survival probability. Hatch marks on survival distributions indicate censored individuals (*i.e*., samples removed for RNAseq sampling or disease testing). Stand-alone survival distributions are presented in Fig. S1 along with family thiamine concentrations and hazard ratio values.

### Gene co-expression network and gene ontology analyses: treatment effect genes

After Bonferroni correction, 3 WGCNA modules of co-expressed genes were significantly correlated with treatment status (corrected *p* < 0.05). Module A contained 667 genes and these genes were associated with 647 significantly overrepresented GO terms; 46 GO terms were unique to Module A and associated with the top genes in the module when genes were ranked by gene significance (terminal nodes in Fig. 3A, Table S4). Many GO terms associated with genes in Module A were related to neurological function and development, including regulation of long-term neuronal synaptic plasticity, neurotransmitter secretion, and neuromuscular junction development (Fig. 3A). Differential expression of genes involved in neurological function may underlie the abnormal locomotion patterns observed in thiamine deficient fry. Module B contained 355 genes associated with 261 significantly overrepresented GO terms; 17 GO terms were unique to Module B and associated with the top genes in the module. Of these 17 GO terms, 8 were associated with metabolism, including positive regulation of insulin secretion, glutamine metabolic process, and tricarboxylic acid metabolic process (Fig. 3B). Differential expression of genes related to these terms is likely related to diminished metabolic rates in untreated individuals. Module C contained 470 genes associated with 768 significantly overrepresented GO terms; 51 GO terms were unique to Module C and associated with the top genes in the module. Many of these GO terms were related to cardiovascular function and development, such as oxygen transport, endocardium formation, and blood vessel maturation (Fig. 3C).

**Figure 3.**
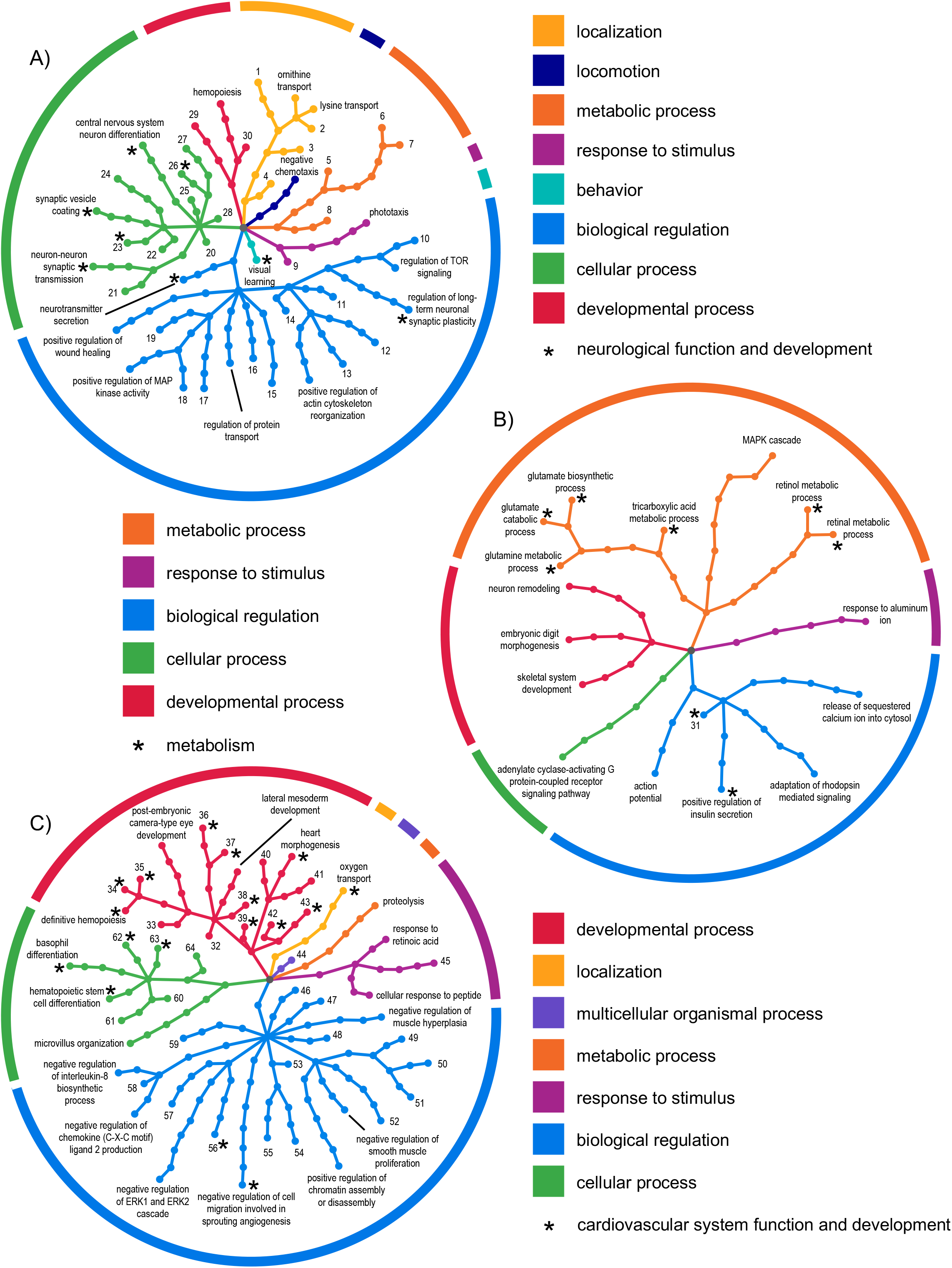
Gene ontology (GO) hierarchy networks constructed using the *metacoder* package in R for three modules (A, B, and C) of co-expressed genes significantly associated with treatment status. Included GO terms were unique to each module and were associated with at least one of the top 20 genes in that module when genes were ranked by WGCNA gene significance. Branch and node colors indicate the biological process child term to which distal nodes belong, with the central grey node representing the biological process level of the GO hierarchy. Terms associated with numbered terminal nodes are provided in Table S4. Terminal nodes marked with an * indicate nodes related to a function or process commonly represented in that module network (*e.g*., several terminal nodes in Module A are related to neurological function and development).

Additionally, all three modules contained terms related to vision, including visual learning, retinal metabolic process, adaptation of rhodopsin mediated signaling, and post-embryonic camera-type eye development. Differential expression of genes related to these terms in untreated individuals is likely associated with decreased visual acuity documented in thiamine deficient fry (Carvalho et al., 2009). Each module also contained DEGs identified through differential expression analysis (representing 23.4%, 18.3%, and 24.5% of genes in each module, respectively). The DEGs assigned to module A were downregulated in treated individuals, while the DEGs assigned to modules B and C were upregulated in treated individuals (Fig. S3).

### Putatively adaptive genes

Maternal effects may influence among-family variation in survival and gene expression. However, we could not identify any maternal characteristics, including maternal thiamine concentrations, that were associated with survival of untreated offspring. Specifically, maternal size and weight were not correlated with untreated offspring survival rate (standard length: F_1,32_ = 0.15, *p* = 0.70; weight: F_1,31_ = 0.10, *p* = 0.75), indicating that differences in survival among families is not simply a function of maternal condition (Fig. 4A,B). Furthermore, for the 9 families sampled for RNA-seq, no relationship appears to exist between maternal muscle or egg thiamine concentrations and proportion of untreated offspring surviving (n = 8 and n = 9, respectively; Fig. 4C,D). Results of linear regressions also indicated that egg thiamine concentration was not a significant predictor of log(hazard ratio) (F_1,7_ = 2.18, *p* = 0.18; Fig. S4). Although we cannot entirely rule out the influence of maternal effects, these results suggest that maternal effects are not driving all of the among-family variation in survival.

**Figure 4.**
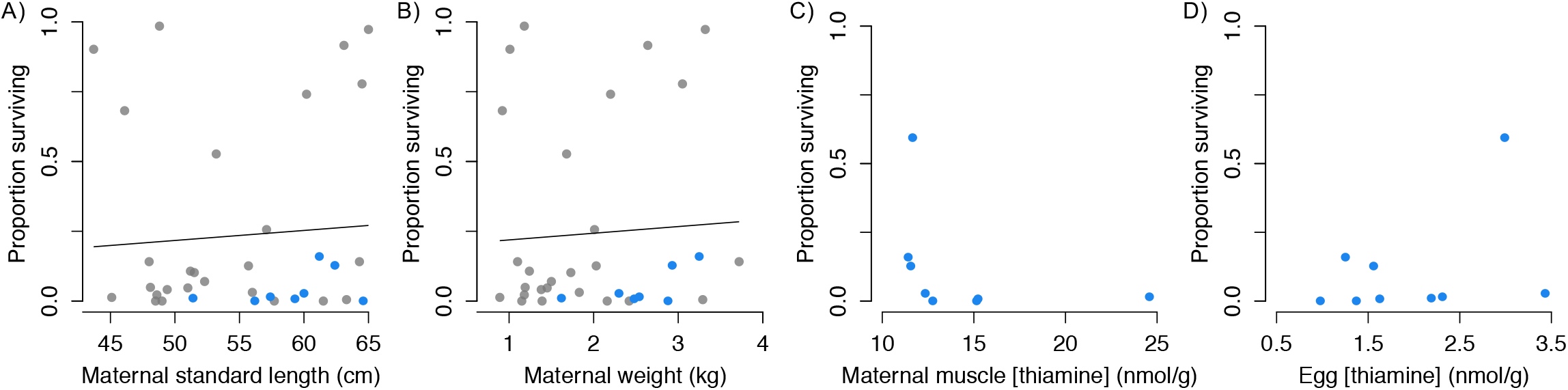
No associations were found between maternal characteristics (weight (kg) and standard length (cm), respectively) and proportion of untreated offspring surviving at the end of the experiment, when tested with linear regressions (A,B). There also does not appear to be any relationship between maternal muscle or egg thiamine concentrations (nmol/g) and proportion of offspring surviving for the 9 families sample for RNA-seq (C,D). In A-D, families sampled for RNA-seq are indicated by blue points; note that data were unavailable for some families in panels A-C (*i.e*., 1 family from panels A and C and 2 families from panel B).

Differential expression analyses conducted using a count matrix for only untreated individuals (n = 18) and with family hazard ratio as the explanatory variable yielded 1,656 DEGs. Of these DEGs, 471 were discarded because the adjusted r^2^ of the regressions for these genes were < 0.3, and an additional 210 were discarded because the regression slopes did not significantly differ from 0. The remaining 1,446 putatively adaptive DEGs were divided into 812 genes positively associated with increased risk of mortality (Fig. 5A,C) and 634 genes negatively associated with increased risk of mortality (Fig 5B,C; Table S5).

**Figure 5.**
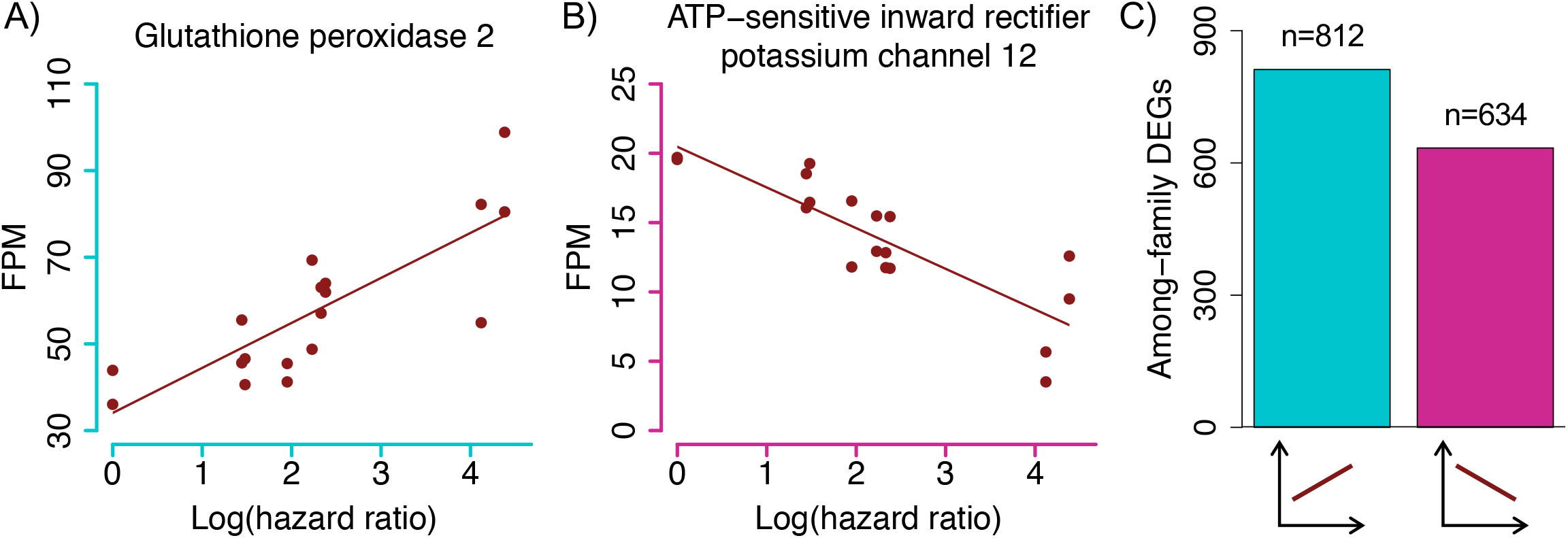
Relative expression of A) glutathione peroxidase (LOC106583190) and B) ATP-sensitive inward rectifier potassium channel 12 (LOC106600689) in fragments per million mapped (FPM) across log(hazard ratio) values. C) The number of putatively adaptive genes positively and negatively associated with increasing risk of mortality. Putatively adaptive genes positively associated with increasing risk of mortality are largely associated with gene ontology terms related to physiological stress, whereas genes negatively associated with risk are associated with terms related to growth and developmental processes (see Table S6).

Adaptively expressed DEGs with positive and negative slopes were associated with 870 and 741 overrepresented GO terms, respectively (*p* < 0.001). Of the top 50 terms associated with genes in each slope category, 17 terms were shared between the categories (Table S6). Shared terms were related to a variety of processes, including regulation of transcription, response to glucose, aging, and oxidation-reduction process. Terms associated with genes with negative slopes (*i.e*., genes upregulated in families with high survival) relate to growth and developmental processes, including cellular proliferation, DNA replication, embryo development, neurogenesis, and visual perception (Table S6). Genes with positive slopes (*i.e*., genes upregulated in families with low survival) were associated with terms that seems to indicate stressful physiological conditions, including response to hydrogen peroxide, response to hypoxia, response to toxic substance, and several terms related to toll-like receptor signaling pathways (Table S6).

### Additive and treatment x among-family effect genes

The differential expression of 114 genes in response to thiamine treatment was driven entirely by additive effects, meaning that the response to treatment was equal among families (Fig. 6A). Of these 114 genes, 84 genes also showed no among-family variation in gene expression, suggesting that the response to thiamine in this group of genes is entirely environmental (*i.e*., not genetic). For 30 additively expressed genes, we also identified significant among-family variation in expression, suggesting that these genes are both putatively adaptive and additive (*i.e*., they respond equally across families) (Fig. 6A,B). For example, expression level of popeye domain-containing protein 2 (*popdc2*; SI Discussion) decreases with increasing hazard ratio rank for both treatment groups, with equal slopes between treatments (Fig. 6B).

**Figure 6.**
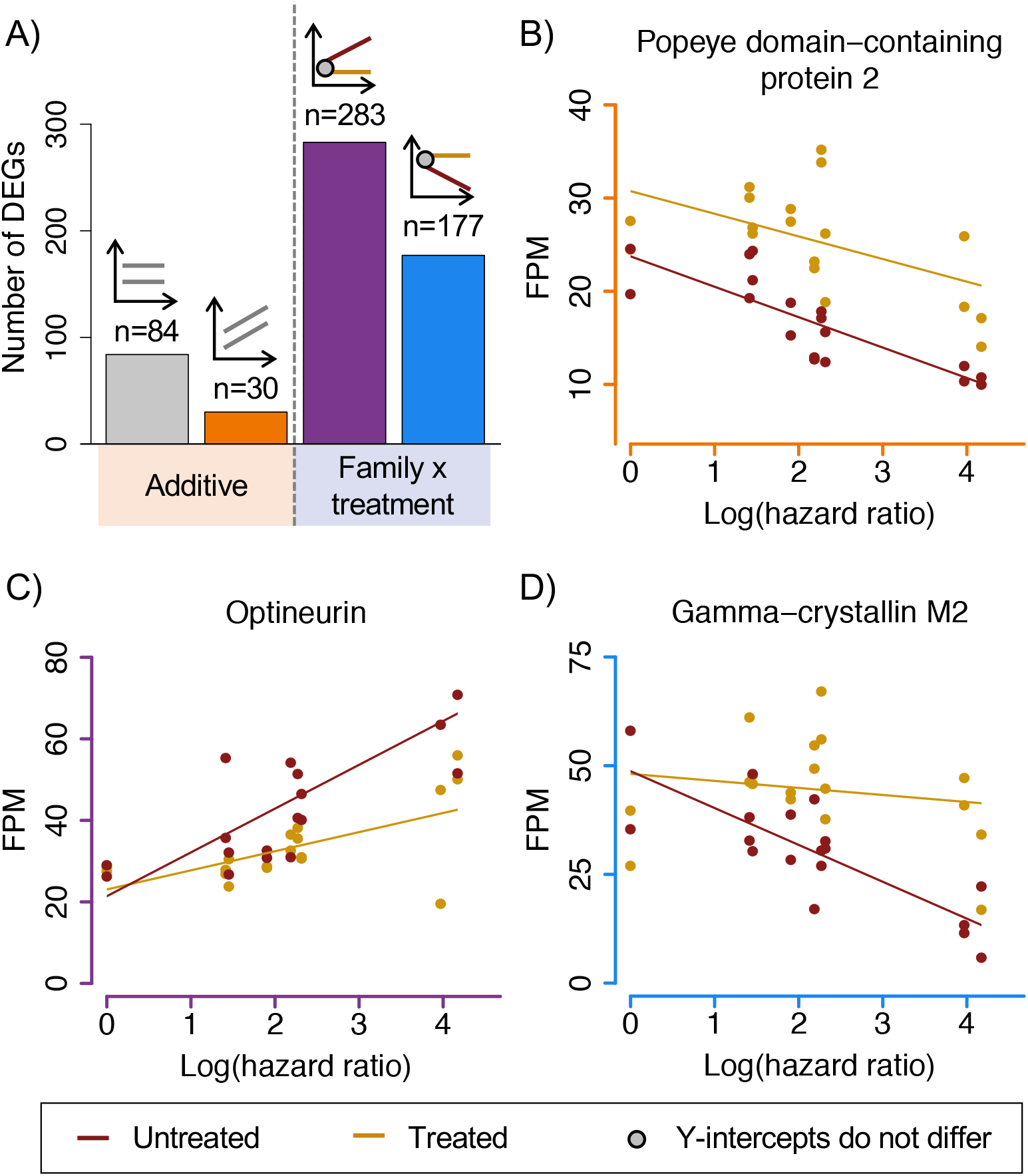
A) The number of genes differentially expressed in response to treatment and among-family differences in 4 categories (from left to right): 1) regression slopes are equal between treatment groups and expression is equal across families, indicating a purely environmental effect of treatment on differential gene expression; 2) regression slopes are equal, indicating that families respond evenly to treatment, but among-family differences suggest putatively adaptive responses; 3) and 4) expression levels are equal between treatment groups in high-surviving families with expression patterns diverging in low-surviving families, indicating putatively adaptive responses. Grey lines in example plots in (A) may represent regression lines for either treated or untreated individuals. B-D) Genes exhibiting expression patterns represented in (A). Axes colors correspond to bar colors in (A). Relative expression of B) popeye domain-containing protein 2 (*popdc2*), C) optineurin (*optn*), and D) gamma-crystallin M2 (LOC106575874) in fragments per million mapped (FPM) across log(hazard ratio) values.

For 597 genes differentially expressed with respect to treatment, the slopes of the regressions for each treatment group differed, indicating a family x treatment effect. The vast majority (460/720) of family- and treatment-effect genes fell into two categories (Fig. 6A), both of which had a shared y-intercept (see Fig. S5 for all identified categories). Gene expression between treated and untreated individuals was most similar at lower hazard ratio values, with expression levels of treated and untreated individuals diverging with increasing hazard ratio. For example, expression levels of the optineurin and gamma-crystallin M2 genes did not differ between treatments for the family with the highest survival (lowest hazard ratio value) (Fig. 6C,D). These two genes differ in their responses to treatment; thiamine treatment decreases expression of optineurin in low survival families, whereas treatment increases expression of gamma-crystallin M2 in low survival families. Because the slopes of the treatment group regressions differ, treatment does not evenly affect gene expression across families; the families with the lowest survival (highest hazard ratio values) experienced the largest shifts in expression levels in response to treatment.

## Discussion

Among all families and across both years, we found a high degree of variation in survival (Fig. 1B) and egg thiamine concentrations, with most females producing eggs that cannot survive without supplemental treatment (Fig. 1C). The ubiquity of low egg thiamine in these samples is consistent with extremely limited reproductive success documented in Lake Champlain tributaries (Prévost, Hill, Grant, Ardren, & Fraser, in press). Treatment with supplemental thiamine does improve survival outcomes for all families, but does not guarantee survival in families with higher hazard ratios (Fig. S1D-I). Egg thiamine concentration could not predict survival (hazard ratio value) for the 9 families included in gene expression analyses (Fig. S4), and no relationships appear to exist between maternal muscle or egg thiamine concentrations and proportion of untreated offspring surviving (Fig. 4C,D). Across all families, maternal length and weight also do not predict offspring survival (proportion surviving; Fig. 4A,B). Because all offspring were raised in a common environment, the absence of this relationship coupled with the high variation in among-family survival indicates that family identity (*i.e*., genetic background) plays an important role in determining whether an individual will survive thiamine deficiency. Furthermore, in the face of thiamine deficiency, certain families are better able to maintain gene expression profiles that approximate expression profiles under thiamine-rich conditions without the aid of supplemental thiamine (Fig. 6), consistent with a genetic basis for tolerance to low thiamine availability.

Across all families, we found that a large number of genes responded to supplemental thiamine treatment. Of the genes hypothesized to be differentially expressed between treated and untreated individuals *a priori*, two gene products perform functions that balance relative intracellular concentrations of thiamine and its various derivatives (see SI text for Discussion; Fig. S2). From the full list of transcript counts, we identified three modules of co-expressed genes associated with treatment status (Fig. 3). For each of these modules, clear themes emerged from their unique lists of overrepresented GO terms. Module A’s association with neurological function and development identified genes related to specific signs of thiamine deficiency in Atlantic salmon fry, such as uncoordinated swimming patterns, inability to maintain an upright position in the water column, and absence of avoidance behavior in response to light exposure (Fisher, Spitsbergen, et al., 1995). These signs of thiamine deficiency may also be related to other overrepresented terms unique to Module A, including responses to stimuli, such as phototaxis and negative chemotaxis. Overrepresented terms in Module B identified genes associated with metabolism, and differential expression of these genes likely underlies slower rates of development under thiamine deficient conditions, with treated individuals achieving larger body sizes than untreated individuals of the same age (Fitzsimons et al., 2009). In Module C, overrepresented terms identified genes related to cardiovascular function and development and may drive vascular dysfunction observed in untreated individuals, as evidenced by hemorrhaging, vascular congestion, and irregular heart rate (Fig. 1A; Fisher, Spitsbergen, et al., 1995). The ubiquity of terms related to vision and eye development shared across all three modules of co-expressed genes demonstrates the complexity of relationships among genes that influence proper development of the visual system (*e.g*., A: visual learning and phototaxis; B: retinol and retinal metabolic processes; C: post-embryonic camera-type eye development).

Differential responses to thiamine deficiency among families comprise 1,446 putatively adaptive genes. These genes are putatively adaptive because their expression level is directly associated with among-family variation in survival. For example, 812 genes are significantly upregulated in untreated individuals from high-survival families (Fig. 5A,C). This result suggests that the increased expression of these genes is associated with higher survival and that these genes, or the various cis or trans acting regulatory elements that influence their expression, could respond to selection in a thiamine-poor environment. Of course, these putatively adaptive genes could also be affected by maternal effects, though survival was not correlated with any maternal or egg traits that we measured (Fig. 4), heritable epigenetic effects (Le Luyer et al., 2017), or other environmental factors. Thus, we are not suggesting that all of these genes would underlie an adaptive response to selection, but rather that this list represents a suite of candidate genes that would likely respond to selection. The fact that there are so many survival-associated genes implicated in an among-family response also suggests that there is sufficient underlying genetic variation in the population to respond to selection. This result, coupled with our survival data, suggest that this population could adaptively respond to selection in the wild. It is worth noting that there is no natural reproduction in Lake Champlain and that all released salmon are treated with supplemental thiamine; this relaxed natural selection could be limiting the successful reintroduction of salmon into the wild.

Of the 3,616 treatment-effect DEGs, 84 displayed evidence of an additive effect of treatment that was not associated with among-family survival (Fig. 6A). In other words, these 84 genes responded to thiamine treatment equally across families and represent a consistent environmental response to the treatment condition. The lack of association between among-family variation in survival and the expression of these genes and the fact that these genes changed in expression roughly equally across families suggests that the change in expression due to thiamine is an entirely environmental response. Thus, we would not predict these genes to respond to selection in a thiamine-poor environment. A small subset of putatively adaptive genes (n = 30) exhibited an additive response to treatment across families (*i.e*., the slopes of the regressions were non-zero but did not differ with respect to treatment) (Fig. 6A,B; SI Discussion). By contrast, 597 DEGs were adaptively expressed and provided evidence of a family x treatment interaction. Most of the genes exhibiting both treatment and family effects (460/720 or 64%) followed predictable patterns of expression that can be broadly grouped into two out of six identified categories (Fig 6D; see Fig. S5 for all family x treatment patterns). For both categories, expression is most similar between treatments in families with high survival and begins to diverge as survival declines. These results illustrate that treatment can have different effects on different families. The fact that the majority (64%) of family x treatment-effect genes occur in the two categories with shared y-intercepts indicates that a successful response to thiamine-poor conditions involves the maintenance of homeostatic conditions.

## Conclusion

Across families, we found no relationship between egg thiamine concentration, maternal traits, and the risk of mortality due to thiamine deficiency, indicating that among-family genetic variation plays an important role in determining thiamine deficiency outcomes. Through gene co-expression network analyses, we determined that many GO terms associated with top DEGs are consistent with observed behavioral and physical signs of thiamine deficiency. Specifically, terms related to neurological function and development, metabolism, cardiovascular function, and visual system development parallel signs of deficiency including uncoordinated swimming patterns, stunted growth, irregular heart rate, and decreased visual acuity. We also described two broad categories of gene expression patterns in response to thiamine deficiency: (1) putatively adaptive genes, which underlie family-level differences in tolerance to low thiamine availability and represent candidate genes likely to respond to selection, and (2) treatment effect genes, which comprise additive and family x treatment effect responses to changes in available thiamine. An additive response coupled with no among-family variation in expression identifies genes that are plastic and respond purely as a function of the treatment condition (i.e., different environments). Such genes would be useful to identify in scenarios where a response to selection was not desirable (e.g., captive breeding programs). Family x treatment effect genes, on the other hand, can be associated with differences in among-family variation in survival, and illustrate that genetic background can differentially affect patterns of gene expression. More importantly, our results identified putatively adaptive genes that would likely respond to selection and that are directly associated with among-family variation in survival. Precisely how much of a response to selection in the wild could occur remains unknown, but uncovering the adaptive genetic variation required for a response to selection represents the first step towards the successful management and conservation species threatened by changing environmental conditions.

## Acknowledgements

We thank T. Chairvolotti and K. Kelsey of the Vermont Fish and Wildlife Department for coordination of spawning and gamete collection events. We also thank H. Bouchard, P. Boynton, S. Frost, T. Jones, E. Lehnert, W. Olmstead, N. Staats, and D. Wong of the US Fish and Wildlife Service for coordination of spawning events, disease testing of adult salmon and offspring, collection of mortality data, rearing of offspring, and assistance with all logistical aspects of this study. Additionally, we thank the Purdue Genomics Core for their sequencing efforts, the Rinchard lab at the State University of New York College at Brockport for thiamine concentration analyses, C. Searle for providing code and assistance with survival analyses, and C. Schraidt and M. Sparks for constructive comments and discussion. This research was funded by the Alton A. Lindsey Graduate Fellowship in Ecology (Purdue University) awarded to AMH and from support from the Purdue Department of Biological Sciences to MRC. The findings and conclusions in the article are those of the authors and do not necessarily represent the views of the U.S. Fish and Wildlife Service.

## Data Accessibility Statement

Upon acceptance, code and scripts will be made available at https://github.com/ChristieLab, and aligned reads (Table S1) will be made available via NCBI Sequence Read Archive with accession numbers provided.

## Author Contributions

AMH, WRA, and MRC designed the project. AMH and WRA collected gametes. AMH performed molecular work. AMH and JRW analyzed data. AMH and MRC wrote the paper. All authors read and approved the final manuscript.

