## Supplemental Information for "Among family variation in survival and gene expression uncovers adaptive genetic variation in a threatened fish"

### Introduction

Coordinated reintroduction efforts for Lake Champlain Atlantic salmon began in 1972 through stocking hatchery reared salmon, but self-sustaining wild populations have not been reestablished. A major limiting factor to the reintroduction effort was the illegal introduction of the non-native alewife to Lake Champlain in 2003 (Marsden & Hauser, 2009). Alewife currently comprise a sufficient proportion of Atlantic salmon diets to cause thiamine-related mortality in eggs collected for hatchery rearing (Marsden & Langdon, 2012), and the Atlantic salmon population remains entirely supported by hatchery supplementation. In the hatchery, fertilized salmon eggs must be treated with thiamine to prevent high rates of mortality due to thiamine deficiency

### Methods

#### *Experimental crosses*

We transported gametes at 4 °C to the White River National Fish Hatchery (Bethel, Vermont, USA), where we systematically combined milt and eggs to generate 35 families (32 males paired to 35 females; in 2016, low availability of males required that one male be crossed with three females and that an additional male be crossed with two females; half-sib families were used in dose-response curve calculations, but not for RNAseq or Cox proportional hazards regressions. To prevent pathogen introduction to the hatchery, we treated all fertilized eggs with a 0.005% iodophor solution for 30 minutes. We next rinsed fertilized eggs with fresh water to prepare them for treatment with thiamine (treated) or water (untreated).

### Discussion

#### *Genes hypothesized to be differentially expression* a priori

One thiamine derivative, thiamine diphosphate (TDP), is required as a cofactor for several enzymes involved in energy production. Conversion of TDP to thiamine triphosphate is catalyzed by adenylate kinase (Fig. S2). Under low thiamine conditions, downregulation of two paralogous genes encoding adenylate kinase may decrease the amount of TDP converted to thiamine triphosphate, resulting in conservation of intracellular TDP pools. Reduced folate carrier transports folate and phosphorylated thiamine derivatives across the cell membrane, and decreased expression of this carrier is associated with increased accumulation of intracellular TDP (Zhao et al., 2001). Downregulation of reduced folate carrier in untreated individuals may also conserve intracellular TDP by reducing the amount of TDP that is exported from the cell. Maintenance of cellular TDP concentration is required for proper cellular metabolism

(Bettendorff, 2013), and adenylate kinase and reduced folate carrier may comprise an important cellular response to thiamine-poor conditions.

The downregulation of thiamine transporter 2 in untreated samples is unexpected, given extensive documentation of increased thiamine transporter 2 mRNA transcription and promoter activity under thiamine deficiency conditions in a variety of study organisms and tissue types (Nabokina, Subramanian, Valle, & Said, 2013; Reidling & Said, 2005). Our observation may conflict with these studies because expression of thiamine transporter 2 is dependent on developmental stage, with expression generally decreasing with maturation (Reidling, Nabokina, Balamurugan, & Said, 2006). Downregulation of thiamine transporter 2 under thiamine deficiency has been described in human cell line culture (Nabokina et al., 2013) and mature mice (Reidling & Said, 2005), but the combined effects of thiamine deficiency and developmental stage on transporter expression have not been evaluated. Juvenile individuals may respond to thiamine deficiency differently than mature individuals at the level of gene expression.

*Popeye domain-containing protein 2: additively expressed and putatively adaptive*

Popeye domain-containing protein 2 (*popdc2*), an additively and adaptively expressed gene, plays a crucial role in cardiac and skeletal muscle development, as evidenced by abnormal muscle fiber morphology, pericardial edema, and irregular heart rate observed in *popdc2*-knockdown zebrafish (*Danio rerio*) (Kirchmaier et al., 2012). These effects of *popdc2*-knockdown are mirrored in thiamine deficient juvenile Atlantic salmon (Essa et al., 2011; Fisher, Spitsbergen, Iamonte, Little, & Delonay, 1995; Sechi & Serra, 2007). Genes exhibiting patterns similar to *popdc2* may represent differences in how thiamine is used or allocated across different genetic backgrounds, regardless of thiamine status (*i.e.*, treated or untreated) (Fig. 6A,B).

*Example family x treatment genes*

Expression of optineurin (*optn*) falls in the first family x treatment category, with thiamine treatment decreasing expression across families with the greatest decrease occurring for families with low survival (Fig. 6C). Accumulation of optineurin has been documented in nervous system tissues affected by such neurodegenerative diseases as amyotrophic lateral sclerosis (ALS), Parkinson's disease, Creutzfeldt-Jakob disease, and glaucoma (Osawa et al., 2011). Another gene following a similar pattern, gamma-crystallin M2, is differentially upregulated in response to thiamine treatment (Fig. 6D). Gamma-crystallin proteins are the primary lens proteins in the vertebrate eye, and high concentrations of these proteins are required for proper lens clarity (H. Zhao, Brown, Magone, & Schuck, 2011). Multiple copies of gamma-crystallin M2 (n = 3) and M3 (n = 6) are differentially downregulated in families with low survival. Downregulation of this gene has previously been described in Atlantic salmon with diet-induced cataracts (Tröße et al., 2009). In our study, downregulation of this gene in families with low survival may lead to gamma-crystallin protein concentrations insufficient for normal lens development, and may help explain diminished foraging efficiency, predator avoidance, and visual acuity in juvenile salmonids that are thiamine deficient (Carvalho et al., 2009; Fitzsimons et al., 2009).

*Example putatively adaptive genes*

For genes with expression levels positively associated with mortality risk (n = 812), related overrepresented GO terms are broadly indicative of physiological stress (Table S6). Genes upregulated in individuals at greater risk of mortality may serve to mitigate unfavorable cellular conditions. For example, one such putatively adaptive gene belongs to the glutathione peroxidase family, a group of enzymes found in all domains of life with antioxidant functions (Toppo, Vanin, Bosello, & Tosatto, 2008). Antioxidant defense mechanisms serve to prevent oxidative stress, which results from reactive oxygen species production and can cause tissue damage (Betteridge, 2000). In our study, expression of the glutathione peroxidase 2 gene increased with decreasing survival in untreated individuals, indicating the need for a stronger response to oxidative stress in families with lower survival rates (Fig. 5A). This result is consistent with previous findings that thiamine deficiency increases glutathione peroxidase expression in Baltic Sea Atlantic salmon (Lundström et al., 1999) and that inactivation of one glutathione peroxidase gene leads to oxidative stress in mice (Esposito et al., 2000). Oxidative stress often accompanies thiamine deficiency (Vuori & Nikinmaa, 2007), and due to the capacity of thiamine to act as an antioxidant, oxidative stress may exacerbate thiamine deficiency, and vice versa (Bettendorff, 2013). This positive feedback loop is observed in many human neurodegenerative disorders, including Alzheimer's disease, Parkinson's disease, and Huntington's disease (Gibson & Zhang, 2002), and may be occurring in families with low rates of survival in our study.

Putatively adaptive genes negatively associated with mortality risk were associated with GO terms related to development and growth (*e.g.*, embryo development, neurogenesis, and kidney development; Table S6). Upregulation of genes in this category likely underlie maintenance of normal cellular processes in families exhibiting higher rates of survival under thiamine-poor conditions. For example, ATP-sensitive inward rectifier potassium channel 12 (*kcnj12*) belongs to a family of potassium channels with important roles in heart rate regulation (Anumonwo & Lopatin, 2010; Vornanen, 2017) (Fig. 5B). In addition to *kcnj12*, two other members of this family (2 copies of *kcnj1* and 2 copies of *kcnj15*) are also differentially expressed across families, with expression upregulated in families exhibiting higher survival.

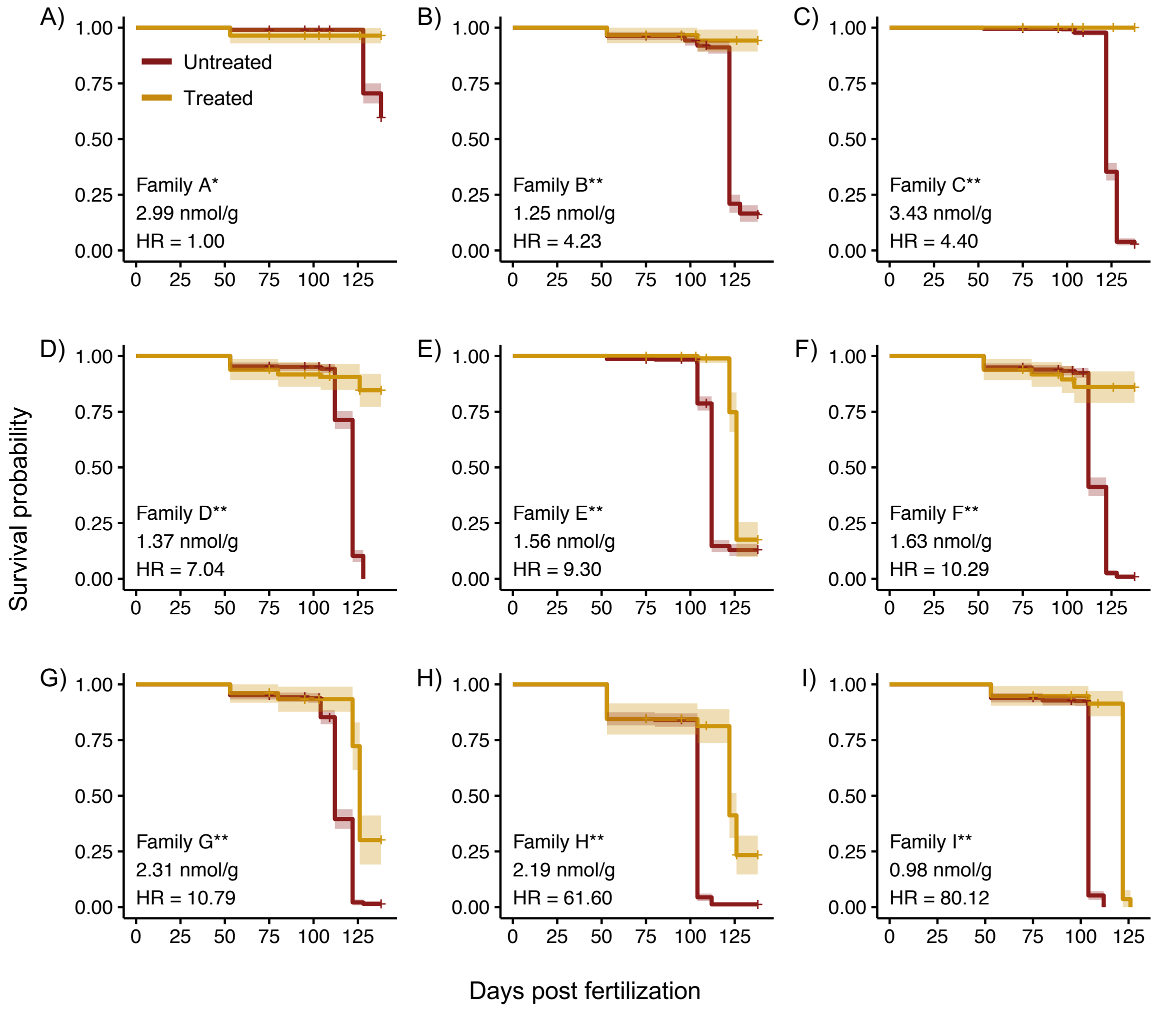


**Figure S1.** Kaplan-Meier survival distributions for treated and untreated groups within each family A-I. Significant differences between treated and untreated survival distributions within each family denoted by * ($p < 0.01$) and **($p < 0.0001$). Total unfertilized egg thiamine concentration for each family is provided, as well as hazard ratios (HR) calculated using a Cox proportional hazards regression. All HR values were calculated using the Family A untreated group as the reference group. Hatch marks on survival distributions indicate censored individuals (*i.e.*, samples removed for RNAseq sampling or disease testing). The eye-up stage was reached at 50 days post-fertilization (dpf) and sampling for RNASeq was conducted at 95 dpf.

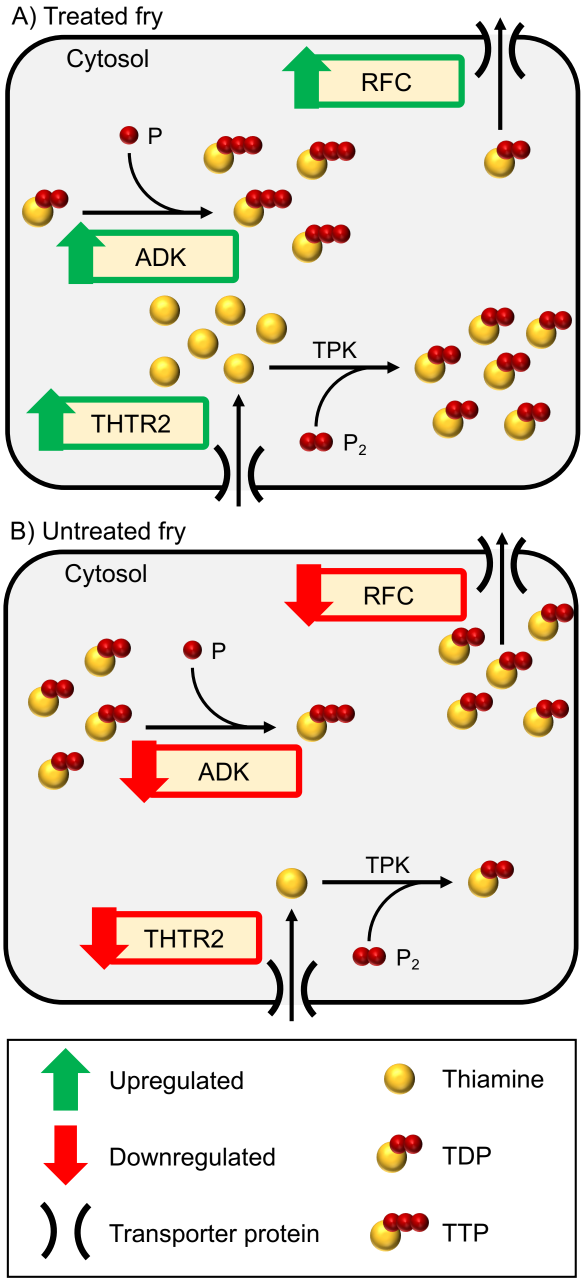


**Figure S2.** Schematic describing how concentrations of thiamine diphosphate (TDP) may be impacted by thiamine status (treated vs. untreated) at the cellular level through differential expression of genes hypothesized *a priori* to be affected by thiamine deficiency. Direction of gene regulation in treated vs. untreated individuals is indicated by arrows next to gene names. THTR transports thiamine across the cell membrane, ADK interconverts forms of thiamine, and RFC exports thiamine derivates from the cell. ADK = adenylate kinase; P = phosphate group; THTR2 = thiamine transporter 2; TPK = thiamine pyrophosphokinase; TTP = thiamine triphosphate; RFC = reduced folate carrier.

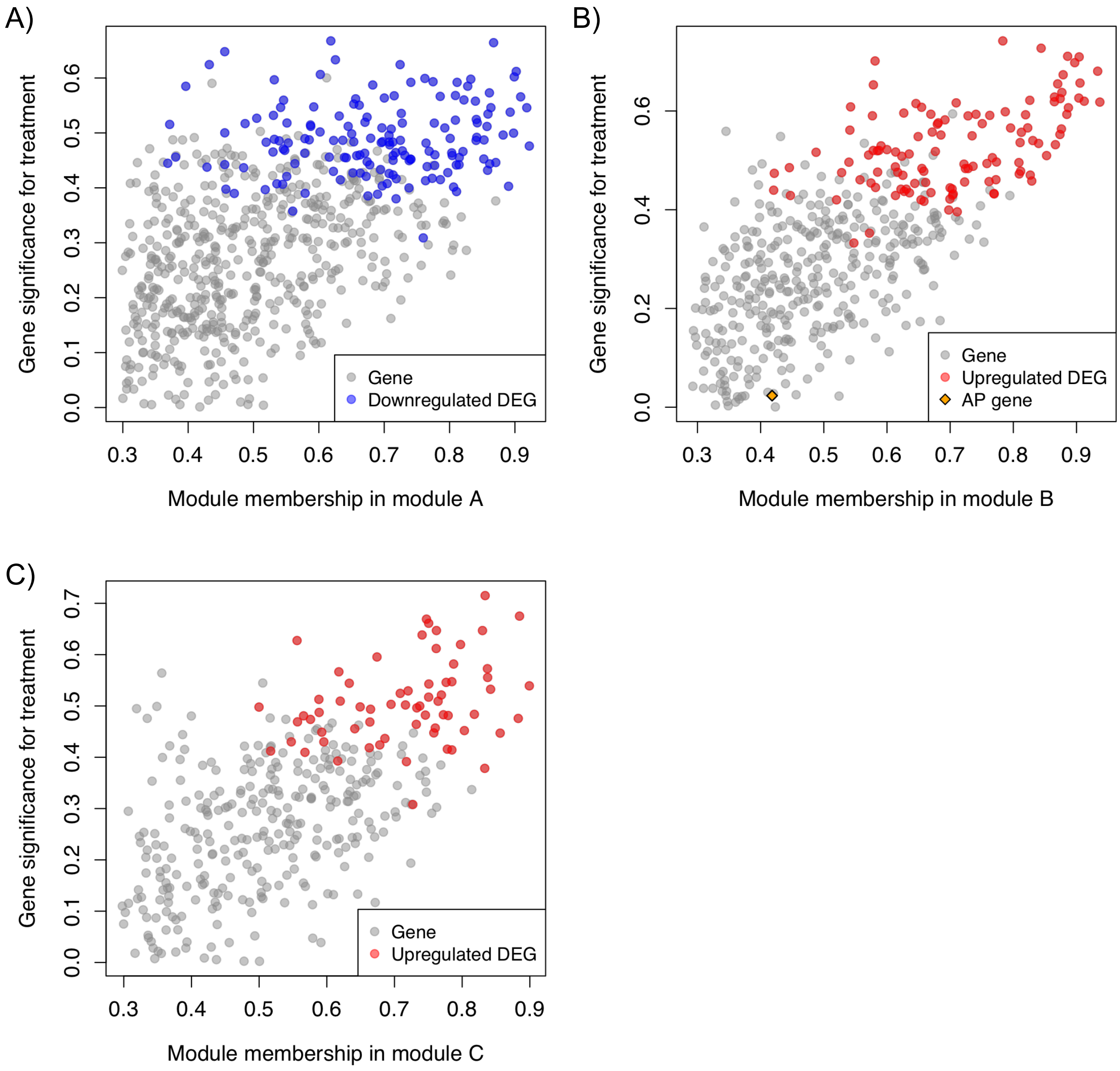


**Figure S3.** Module membership vs. gene significance for each of the modules (A-C) that were significantly correlated with treatment status after Bonferroni correction. Module membership is a value calculated during WGCNA module construction that indicates how many connections (*i.e.*, how tightly correlated in terms of expression) a gene has to other genes within that module. Genes that are significantly differentially expressed are highlighted in blue (downregulated) or red (upregulated). Genes hypothesized to be differentially expressed *a priori* (AP) are indicated by an orange diamond. DEG = differentially expressed gene.

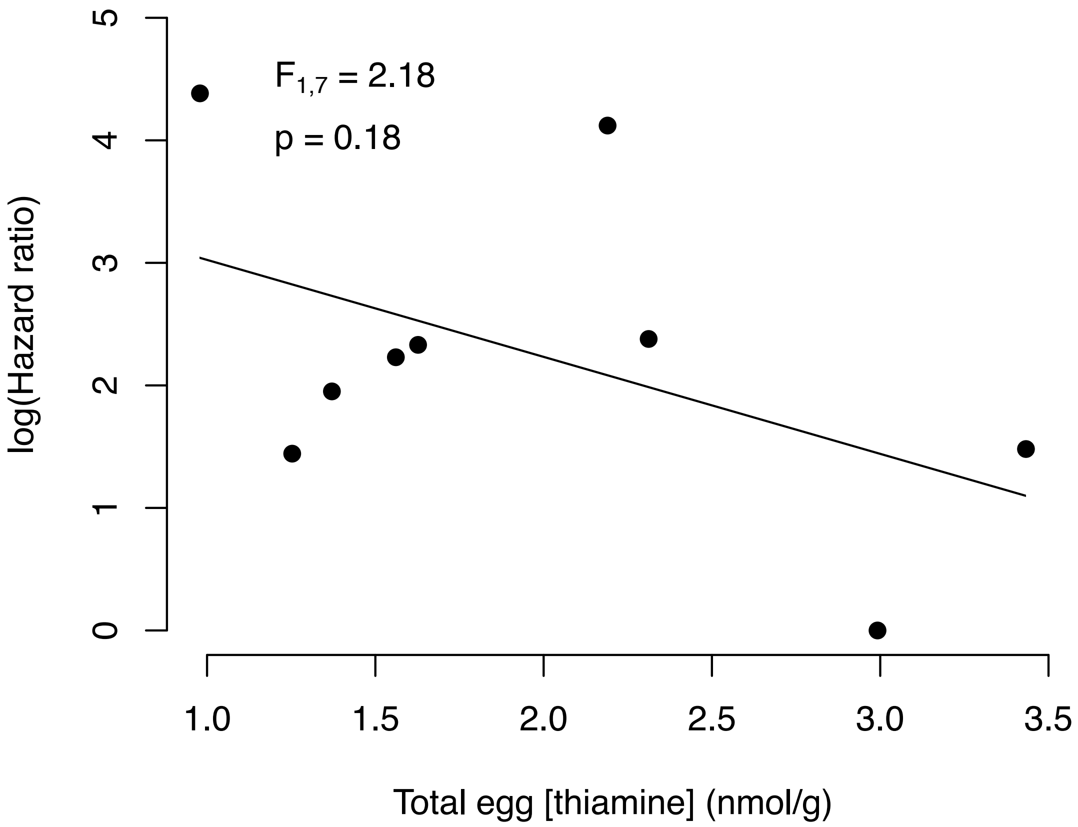


**Figure S4.** Linear regression results demonstrating no relationship between total egg thiamine concentration (nmol/g) and hazard ratio ($\log_e$). This pattern indicates that individual survival is not simply a product of egg thiamine allocation.

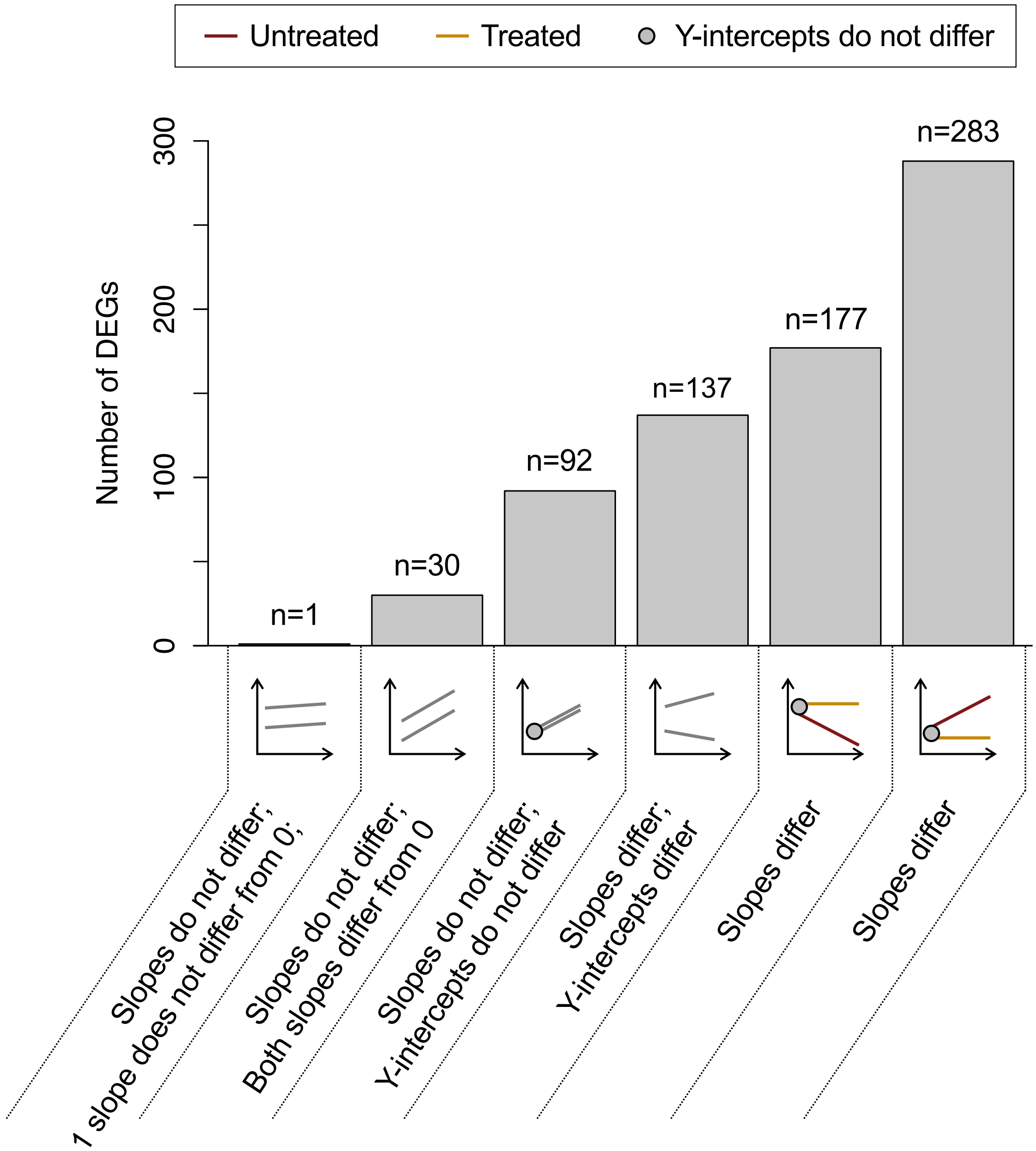


**Figure S5.** Six categories into which all family x treatment differentially expressed genes (DEGs) can be classified (n = 720). Grey lines can represent either treated or untreated regression lines. Numbers of DEGs in each category are provided, as well as an example plots and statistical description for each category. For example plots below the *x*-axis of the barplot, the *x*-axis is log(hazard ratio) and the *y*-axis is fragments per million mapped (FPM). These example plots are representative of the criteria used to define each category. Empirical examples of these types of plots may be found in Fig. 5A,B and Fig. 6B-D in the main text. Although all 720 genes represented here are family x treatment DEGs, our stringent filtering of regression results identified 92 genes as having slopes and *y*-intercepts that do not differ despite being identified as differentially expressed with DESeq2.

**Table S1.** Number of pairs of reads generated per individual (*i.e.*, sequencing library). Sample names correspond to filenames of aligned reads available on NCBI GenBank as BAM files (Accession # will be provided here). Treatment status is provided, with "+" and "-" corresponding to thiamine treated and untreated individuals, respectively.

| Sample name | Number of read pairs | Family | Treatment |
|---|---|---|---|
| 13-a | 41,288,834 | A | - |
| 13-b | 41,577,084 | A | - |
| 13a2 | 38,865,181 | A | + |
| 13b | 26,864,961 | A | + |
| 1-a | 35,780,209 | B | - |
| 1-b | 42,711,237 | B | - |
| 1b | 40,088,378 | B | + |
| 1c | 33,165,177 | B | + |
| 8-a | 50,995,983 | C | - |
| 8-b | 37,425,847 | C | - |
| 8a | 51,873,841 | C | + |
| 8b | 55,729,099 | C | + |
| 9-a2 | 32,770,553 | D | - |
| 9-b | 29,347,029 | D | - |
| 9a | 45,354,403 | D | + |
| 9b | 58,629,131 | D | + |
| 11-a | 40,153,476 | E | - |
| 11-b | 30,827,911 | E | - |
| 11a | 32,286,333 | E | + |
| 11b | 46,616,361 | E | + |
| 3-a | 29,675,435 | F | - |
| 3-b | 49,240,974 | F | - |
| 3a | 42,976,636 | F | + |
| 3b | 35,416,050 | F | + |
| 4-a | 31,610,722 | G | - |
| 4-b2 | 48,885,987 | G | - |
| 4a2 | 28,871,500 | G | + |
| 4b | 37,349,044 | G | + |
| 6-a | 47,444,314 | H | - |
| 6-b | 42,169,359 | H | - |
| 6a | 33,180,043 | H | + |
| 6b | 33,761,100 | H | + |
| 7-a | 30,420,203 | I | - |
| 7-b | 30,604,518 | I | - |
| 7a | 41,586,010 | I | + |
| 7b | 40,992,778 | I | + |

**Table S2.** Gene ontology (GO) terms found to be overrepresented in all 3 modules of co-expressed genes.

| GO ID | GO term |
|---|---|
| GO:0030036 | actin cytoskeleton organization |
| GO:0001525 | angiogenesis |
| GO:0048513 | animal organ development |
| GO:0006915 | apoptotic process |
| GO:0007411 | axon guidance |
| GO:0007420 | brain development |
| GO:0007154 | cell communication |
| GO:0008283 | cell proliferation |
| GO:0007267 | cell-cell signaling |
| GO:0006974 | cellular response to DNA damage stimulus |
| GO:0071260 | cellular response to mechanical stimulus |
| GO:0007417 | central nervous system development |
| GO:0007268 | chemical synaptic transmission |
| GO:0000910 | cytokinesis |
| GO:0048813 | dendrite morphogenesis |
| GO:0009792 | embryo development ending in birth or egg hatching |
| GO:0008543 | fibroblast growth factor receptor signaling pathway |
| GO:0030900 | forebrain development |
| GO:0007507 | heart development |
| GO:0006955 | immune response |
| GO:0006954 | inflammatory response |
| GO:0045087 | innate immune response |
| GO:0001889 | liver development |
| GO:0007613 | memory |
| GO:0008045 | motor neuron axon guidance |
| GO:0035264 | multicellular organism growth |
| GO:0043066 | negative regulation of apoptotic process |
| GO:0008285 | negative regulation of cell proliferation |
| GO:0040015 | negative regulation of multicellular organism growth |
| GO:0000122 | negative regulation of transcription by RNA polymerase II |
| GO:0045892 | negative regulation of transcription, DNA-templated |
| GO:0045071 | negative regulation of viral genome replication |
| GO:0001843 | neural tube closure |
| GO:0001764 | neuron migration |
| GO:0007422 | peripheral nervous system development |
| GO:0048518 | positive regulation of biological process |
| GO:0030307 | positive regulation of cell growth |
| GO:0030335 | positive regulation of cell migration |
| GO:0008284 | positive regulation of cell proliferation |
| GO:0051091 | positive regulation of DNA-binding transcription factor activity |
| GO:0048146 | positive regulation of fibroblast proliferation |
| GO:0043123 | positive regulation of I-kappaB kinase/NF-kappaB signaling |
| GO:0043525 | positive regulation of neuron apoptotic process |

| | |
|---|---|
| GO:0010976 | positive regulation of neuron projection development |
| GO:0045860 | positive regulation of protein kinase activity |
| GO:0001934 | positive regulation of protein phosphorylation |
| GO:0048661 | positive regulation of smooth muscle cell proliferation |
| GO:0045944 | positive regulation of transcription by RNA polymerase II |
| GO:0009791 | post-embryonic development |
| GO:0046777 | protein autophosphorylation |
| GO:0006468 | protein phosphorylation |
| GO:0006898 | receptor-mediated endocytosis |
| GO:0042981 | regulation of apoptotic process |
| GO:0065008 | regulation of biological quality |
| GO:0008360 | regulation of cell shape |
| GO:0050794 | regulation of cellular process |
| GO:0006357 | regulation of transcription by RNA polymerase II |
| GO:0006355 | regulation of transcription, DNA-templated |
| GO:0042493 | response to drug |
| GO:0009749 | response to glucose |
| GO:0001666 | response to hypoxia |
| GO:0032496 | response to lipopolysaccharide |
| GO:0014070 | response to organic cyclic compound |
| GO:0010033 | response to organic substance |
| GO:0007165 | signal transduction |
| GO:0007283 | spermatogenesis |
| GO:0007601 | visual perception |

**Table S4.** Gene ontology (GO) terms associated with numbered terminal nodes of GO hierarchy trees in Figure 3. Terms marked with * are related to neurological function and development (Module A), metabolism (Module B), and cardiovascular system development and function (Module C). The central node in all networks represents the biological process level of the GO hierarchy.

| Module | Number | GO term |
|---|---|---|
| A | 1 | amino acid transmembrane transport |
| A | 2 | arginine transport |
| A | 3 | endosomal transport |
| A | 4 | protein localization |
| A | 5 | phosphorylation |
| A | 6 | cAMP catabolic process |
| A | 7 | cGMP catabolic process |
| A | 8 | glycerol ether metabolic process |
| A | 9 | response to endogenous stimulus |
| A | 10 | positive regulation of Rac protein signal transduction |
| A | 11 | positive regulation of necrotic cell death |
| A | 12 | positive regulation of ruffle assembly |
| A | 13 | negative regulation of receptor-mediated endocytosis |
| A | 14 | regulation of cellular senescence |
| A | 15 | negative regulation of leukocyte chemotaxis |
| A | 16 | positive regulation of angiogenesis |
| A | 17 | positive regulation of miRNA metabolic process |
| A | 18 | activation of MAPKK activity |
| A | 19 | positive regulation of gene expression |
| A | 20 | establishment or maintenance of cell polarity |
| A | 21 | lateral inhibition |
| A | 22 | actin filament organization |
| A | 23 | neuromuscular junction development* |
| A | 24 | protein heterooligomerization |
| A | 25 | cell redox homeostasis |
| A | 26 | neuropeptide signaling pathway* |
| A | 27 | Ras protein signal transduction |
| A | 28 | cell death |
| A | 29 | cellular senescence |
| A | 30 | tube development |
| B | 31 | regulation of metabolic process* |
| C | 32 | midbrain development |
| C | 33 | thyroid gland development |
| C | 34 | embryonic hemopoiesis* |
| C | 35 | post-embryonic hemopoiesis* |
| C | 36 | nucleate erythrocyte development* |
| C | 37 | megakaryocyte development* |
| C | 38 | embryonic heart tube development* |
| C | 39 | endocardium formation* |
| C | 40 | cell morphogenesis |

| | | |
|---|---|---|
| C | 41 | epidermis morphogenesis |
| C | 42 | erythrocyte maturation* |
| C | 43 | blood vessel maturation* |
| C | 44 | stem cell population maintenance |
| C | 45 | cellular response to cycloheximide |
| C | 46 | negative regulation of response to cytokine stimulus |
| C | 47 | positive regulation of cell division |
| C | 48 | positive regulation of protein complex assembly |
| C | 49 | regulation of rhodopsin mediated signaling pathway |
| C | 50 | negative regulation of phosphatidylinositol 3-kinase signaling |
| C | 51 | negative regulation of protein kinase B signaling |
| C | 52 | negative regulation of cysteine-type endopeptidase activity involved in apoptotic process |
| C | 53 | regulation of stem cell population maintenance |
| C | 54 | positive regulation of endothelial cell differentiation |
| C | 55 | regulation of mast cell differentiation |
| C | 56 | positive regulation of hemoglobin biosynthetic process* |
| C | 57 | regulation of phosphatidylinositol 3-kinase activity |
| C | 58 | negative regulation of interleukin-6 production |
| C | 59 | negative regulation of heterotypic cell-cell adhesion |
| C | 60 | epidermal cell differentiation |
| C | 61 | type I pneumocyte differentiation |
| C | 62 | hemangioblast cell differentiation* |
| C | 63 | platelet formation* |
| C | 64 | mesodermal cell fate determination |

**Table S6.** Top 50 biological process gene ontology (GO) terms for each category (positive and negative slopes) of adaptively expressed genes, when terms are ranked by *p* value.

| GO ID | GO term | Slope category |
|---|---|---|
| GO:0007568 | aging | both |
| GO:0043066 | negative regulation of apoptotic process | both |
| GO:0008285 | negative regulation of cell proliferation | both |
| GO:0000122 | negative regulation of transcription by RNA polymerase II | both |
| GO:0045892 | negative regulation of transcription, DNA-templated | both |
| GO:0055114 | oxidation-reduction process | both |
| GO:0045944 | positive regulation of transcription by RNA polymerase II | both |
| GO:0050794 | regulation of cellular process | both |
| GO:0006355 | regulation of transcription, DNA-templated | both |
| GO:0051591 | response to cAMP | both |
| GO:0042493 | response to drug | both |
| GO:0032355 | response to estradiol | both |
| GO:0009749 | response to glucose | both |
| GO:0050896 | response to stimulus | both |
| GO:0007165 | signal transduction | both |
| GO:0044281 | small molecule metabolic process | both |
| GO:0007283 | spermatogenesis | both |
| GO:0031100 | animal organ regeneration | negative |
| GO:0008283 | cell proliferation | negative |
| GO:0007166 | cell surface receptor signaling pathway | negative |
| GO:0044260 | cellular macromolecule metabolic process | negative |
| GO:0009987 | cellular process | negative |
| GO:0006974 | cellular response to DNA damage stimulus | negative |
| GO:0007268 | chemical synaptic transmission | negative |
| GO:0006260 | DNA replication | negative |
| GO:0006271 | DNA strand elongation involved in DNA replication | negative |
| GO:0006268 | DNA unwinding involved in DNA replication | negative |
| GO:0009790 | embryo development | negative |
| GO:0007588 | excretion | negative |
| GO:0007186 | G protein-coupled receptor signaling pathway | negative |
| GO:0000082 | G1/S transition of mitotic cell cycle | negative |
| GO:0001701 | in utero embryonic development | negative |
| GO:0001822 | kidney development | negative |
| GO:0070309 | lens fiber cell morphogenesis | negative |
| GO:0043433 | negative regulation of DNA-binding transcription factor activity | negative |
| GO:0022008 | neurogenesis | negative |
| GO:0043065 | positive regulation of apoptotic process | negative |
| GO:0008284 | positive regulation of cell proliferation | negative |
| GO:0006267 | pre-replicative complex assembly involved in nuclear cell cycle DNA replication | negative |
| GO:0051260 | protein homooligomerization | negative |
| GO:0051289 | protein homotetramerization | negative |

| | | |
|---|---|---|
| GO:0050789 | regulation of biological process | negative |
| GO:0030174 | regulation of DNA-dependent DNA replication initiation | negative |
| GO:0006357 | regulation of transcription by RNA polymerase II | negative |
| GO:0000083 | regulation of transcription involved in G1/S transition of mitotic cell cycle | negative |
| GO:0033993 | response to lipid | negative |
| GO:0035864 | response to potassium ion | negative |
| GO:0006950 | response to stress | negative |
| GO:0035725 | sodium ion transmembrane transport | negative |
| GO:0007601 | visual perception | negative |
| GO:0006915 | apoptotic process | positive |
| GO:0007596 | blood coagulation | positive |
| GO:0007420 | brain development | positive |
| GO:0007623 | circadian rhythm | positive |
| GO:0009792 | embryo development ending in birth or egg hatching | positive |
| GO:0007173 | epidermal growth factor receptor signaling pathway | positive |
| GO:0008286 | insulin receptor signaling pathway | positive |
| GO:0030216 | keratinocyte differentiation | positive |
| GO:0050900 | leukocyte migration | positive |
| GO:0002009 | morphogenesis of an epithelium | positive |
| GO:0006936 | muscle contraction | positive |
| GO:0002755 | MyD88-dependent toll-like receptor signaling pathway | positive |
| GO:0006469 | negative regulation of protein kinase activity | positive |
| GO:0002119 | nematode larval development | positive |
| GO:0048011 | neurotrophin TRK receptor signaling pathway | positive |
| GO:0040010 | positive regulation of growth rate | positive |
| GO:0042127 | regulation of cell proliferation | positive |
| GO:0000003 | reproduction | positive |
| GO:0014823 | response to activity | positive |
| GO:0051384 | response to glucocorticoid | positive |
| GO:0009408 | response to heat | positive |
| GO:0042542 | response to hydrogen peroxide | positive |
| GO:0001666 | response to hypoxia | positive |
| GO:0032496 | response to lipopolysaccharide | positive |
| GO:0009612 | response to mechanical stimulus | positive |
| GO:0032570 | response to progesterone | positive |
| GO:0009636 | response to toxic substance | positive |
| GO:0035914 | skeletal muscle cell differentiation | positive |
| GO:0034134 | toll-like receptor 2 signaling pathway | positive |
| GO:0034138 | toll-like receptor 3 signaling pathway | positive |
| GO:0034142 | toll-like receptor 4 signaling pathway | positive |
| GO:0034146 | toll-like receptor 5 signaling pathway | positive |
| GO:0006418 | tRNA aminoacylation for protein translation | positive |
